# Axon onset remodeling in response to network activity in mouse cortical neurons

**DOI:** 10.1101/2023.07.31.551236

**Authors:** Nadja Lehmann, Stefan Markovic, Christian Thome, Maren Engelhardt

**Affiliations:** Institute of Neuroanatomy, Mannheim Center for Translational Neuroscience (MCTN), Medical Faculty Mannheim, Heidelberg University, Germany; Institute of Anatomy and Cell Biology, Medical Faculty, Johannes Kepler University Linz, Austria; Institute for Stem Cell Biology and Regenerative Medicine, Stanford University, USA

**Keywords:** Axon-carrying dendrite (AcD), axon initial segment, somatosensory cortex barrel field, morphological plasticity

## Abstract

In certain subtypes of pyramidal neurons, axons originate from basal dendrites, resulting in an axon-carrying dendrite branch with unique functional characteristics (AcD cells). The number of AcD cells increases during development, however, it is so far unclear whether neurons remodel their axon emergence throughout their lifetime in response to changes in network activity. To test our hypothesis of such large-scale morphological plasticity, we utilized *in vitro* and *in vivo* strategies in mouse primary somatosensory cortex to test whether network activity impacts axon onset. Based on data obtained by immunofluorescence, confocal microscopy and/or live-cell imaging, we show that neurons are indeed capable of changing the onset of their axon origin from somatic to dendritic and *vice versa* within a few days *in vitro* and that peripheral whisker manipulation and thus changes in sensory input drives large-scale morphological plasticity *in vivo*.

## Introduction

The anatomical diversity of axon origins across neuronal cell populations, in combination with the heterogeneity of length and position of the axon initial segment (AIS), the site of action potential generation, substantially influence the electrical behavior of neurons (reviewed in ^1^). In the traditional view of neuronal morphology, cortical principal neurons feature axons with a somatic onset. Yet, in a subtype of neurons, axons originate from basal dendrites, resulting in an axon-carrying dendrite branch (AcD). This morphology was first observed by Ramon y Cajal in 1937 and since then has been described in numerous cell types, brain areas across species, and model systems ^2-11^. Inputs at these branches have a high likelihood of generating dendritic sodium spikes, triggering action potentials with lower current thresholds, and are partially shielded from perisomatic inhibition ^12^.

A recent study demonstrated that hippocampal CA1 neurons with AcD morphology (AcD cells) are more likely to participate in sharp wave-ripple oscillations, which has important implications for the neuronal correlate of memory consolidation and retrieval ^13^. While AcD cells make up approximately 50% of CA1 pyramidal cells, this trait is also seen in the primary somatosensory cortex, where these cells constitute approximately 30% of thick-tufted layer V neurons ^14^.

Previous observations further showed that the number of AcD cells increases between P8 and P28 in mice ^12^, however, it is so far unclear whether AcD morphology is determined by developmental factors and whether it remains stable throughout the life of a neuron. Considering the various modes of axonal plasticity throughout a neuron’s lifetime and its impact on neuronal network function particularly in the context of the AIS ^15-21^, we hypothesized that neurons may be able to change the position of the axonal arbor between somatic and dendritic in response to changes in network state at time-points past initial developmental phases. If so, such a mode of whole cell plasticity may privilege certain dendritic input channels over others in a dynamic manner and contribute to neuronal plasticity and network adaption.

In the present study, using immunofluorescence and live-cell imaging in organotypic slice cultures (OTC) derived from Thy1-GFP mice, we observed a large-scale remodeling of the entire basal cell pole, occurring both spontaneously and after network manipulation. We further tested axon onset plasticity *in vivo*, using an established sensory deprivation and stimulation paradigm in layer V primary somatosensory cortex, barrel field (S1BF) neurons ^17^. Our data suggest that the position of axon onset is plastic and responds to changes in neuronal activity, thereby potentially complementing synaptic plasticity and neuronal homeostasis.

## Results

### AcD and nonAcD cells exhibit adaptive morphologies

We prepared cortical organotypic slice cultures (OTC) from mice expressing the fluorescent protein GFP in a sparse subset of principal pyramidal cells (Thy1-GFP mouse line). Live imaging of neuronal morphology was performed daily, from DIV3 to DIV10, to capture potential morphological changes (Fig.1A-B). Subsequent immunolabeling of the AIS using a marker against the AIS scaffold protein βIV-spectrin allowed us to separate the axonal branch from dendritic processes (Fig. 1C).

**Figure 1:**
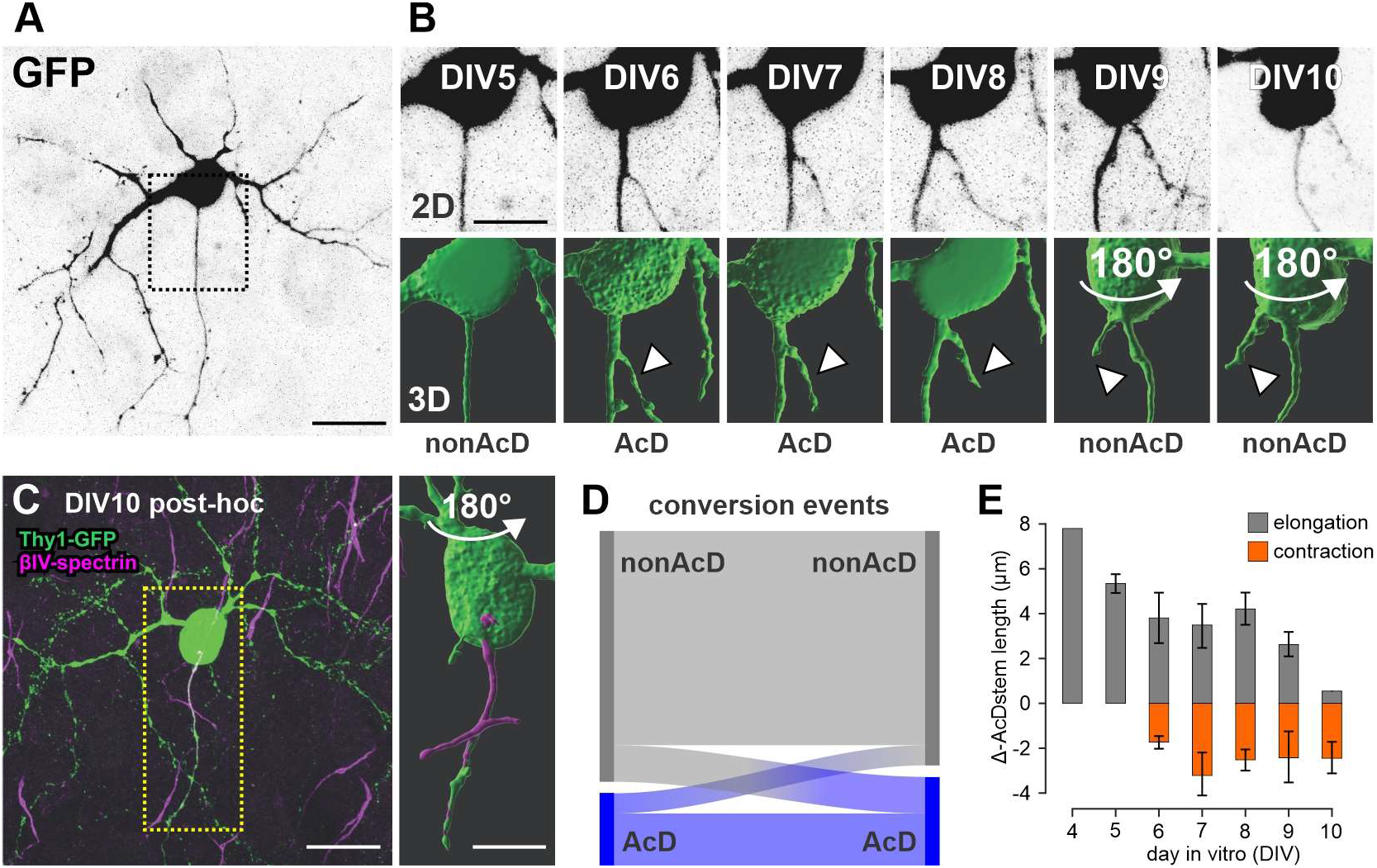
Live-cell imaging demonstrates transitions in axon onset between somatic and dendritic position. (A) Overview of live-imaged Thy-GFP-positive neuron at DIV5. Scale bar = 25 µm. (B) Conversion of an initial nonAcD neuron to an AcD neuron. Close-ups depict the same live-imaged neuron, as shown in B and 3D reconstruction for 6 consecutive days. At DIV6, a dendritic branch (arrowhead) sprouts from the axonal arbor and separates again on DIV9. Scale bars = 12.5 µm (C) Cell from B and C after fixation and immunolabeling of the AIS (βIV-spectrin, magenta). Scale bars = 25 µm (left) and 12.5 µm (right). (D) Sankey diagram visualizing the observed transitions between nonAcD (gray) and AcD morphology (blue) of 98 cells. There were more conversions from nonAcD to AcD state (indicated by the line thickness). Overall, most of the live-imaged neurons retained their axon morphology classification. (E) Barplots depict changes in AcD stem dendrite length observed between imaging sessions from DIV5 to DIV10. One-way ANOVA found no significant differences between elongation and contraction values (n = 17 cells).

Most neurons retained their morphological type (65 nonAcD, 16 AcD of 98 cells imaged for at least two consecutive days). However, in some neurons, the axon onset shifted between a dendritic and a somatic origin, and *vice versa* in a matter of days (total 17 of 98 cells, Fig. 1D). The transformation from a nonAcD to an AcD neuron was observed twice as often compared to AcD to nonAcD (11 vs 6 cells). In one example, we observed two transformations, from nonAcD to AcD and back to nonAcD (example given in Fig.1A-C). In total, we observed five different transformation types: three for conversion to AcD and two for conversion to nonAcD morphology. First, the sprouting of a dendrite from the axonal shaft (observed within 24 h, Fig. 1A; 2 of 11 cases) converts a formerly somatic axon onset into a dendritic one, thus producing AcD morphology. Second, the dendritic arbor and axon converged until they merged into a single branch (Supplement Fig. S1; 2 of 11 cases). Third, the axon relocated from the soma towards the dendrite (Supplement Fig. S2; 6 of 11 cases). The transition from AcD to nonAcD morphology required the retraction of the AcD stem dendrite until the axonal and dendritic branches emerged from the somatic envelope (Fig. 1B; 5 of 6 cases). In one case, we observed that the dendrite gradually separated from the axon until it was sequestered and emanated more laterally from the soma (Supplement Fig. S2).

Based on these observations, we conclude that the axon onset seems to be a dynamic property with the capability to change from AcD to nonAcD and *vice versa* over time. These changes are likely not permanent, as the proximal anatomy of the observed neurons remained dynamic and at least one neuron returned to its original nonAcD morphology.

### Activity-dependent AcD development *in vitro*

Numerous studies demonstrated structural AIS plasticity (length, position within the axon) in response to changes in network activity *in vitro* and *in vivo* ^15-21^. Here we investigated whether such network changes also affect the transition between AcD and nonAcD morphologies in our OTC. We used established *in vitro* pharmacological paradigms to either increase (bicuculline, KCl) or decrease (ifenprodil, MgSO_4_) network activity. Cortical OTC were prepared from P5 animals and treated with one of the above-mentioned reagents for seven days from DIV3 to DIV10 with media changes every second day (Fig. 2A). At DIV10, OTC were fixed and processed for immunofluorescence. Treated and untreated cultures were derived from the alternating hemispheres of the same brain slice to exclude potential hemispherical effects (Fig. 2A).

**Figure 2:**
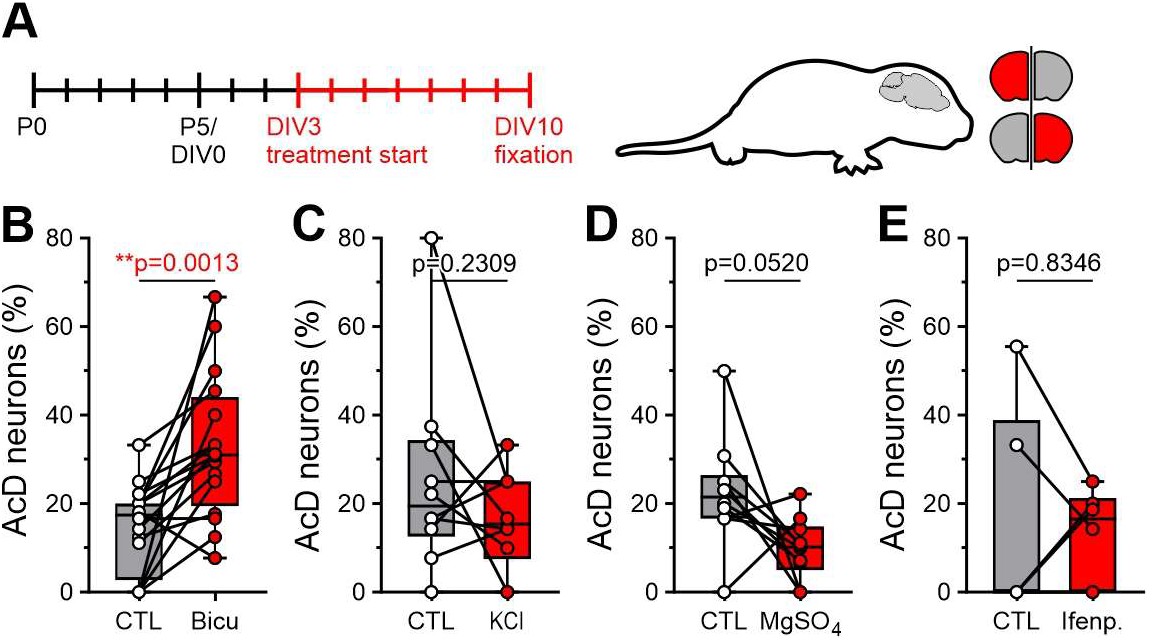
Pharmacological modulators of network activity change the ratio of AcD to nonAcD neurons *in vitro*. (A) Experimental design: OTC were prepared from P5 pups. Treatments started at DIV3 and lasted for 7 days, with media changes every second day (medium + either 6 mM KCl, 15 µM bicuculline, 1 µM ifenprodil hemitartrate, or 10 mM MgSO_4_). Pairs of treated (red) and untreated OTC (gray) were generated from hemispheres of the same brain slice in an alternating manner with the contralateral hemisphere serving as the control. (B) Incubation of OTC with the GABA blocker bicuculline resulted in a significant increase in the number of AcD neurons in the OTC relative to that in the control hemisphere (paired *t*-test, *P*=0.0013, *n*=16). (C) The addition of extracellular KCl, which depolarizes somatic membrane potentials, did not affect the number of AcD neurons (paired *t*-test, *n*=10). (D) Incubation with MgSO_4_ led to fewer AcD neurons compared to the control hemisphere (paired *t*-test, *n*=10), although the change was not statistically significant. (E) Incubation with ifenprodil, a blocker of presynaptic NMDA receptors, did not affect the number of AcD neurons (paired *t*-test, *n*=6).

Bicuculline blocks GABAergic synapses and is commonly used to increase network activity in OTC ^22^. In this study, 15 µM bicuculline was added to the culture medium. Our analysis revealed that the number of AcD neurons in the bicuculline-treated hemisphere was significantly increased compared to the control hemisphere (Fig. 2B, bicuculline-treated OTC: 32.9% ± 16.4% vs. control: 14.6% ± 10.1%; P=0.0013, paired *t*-test, n=16). However, when we used an elevated extracellular KCl concentration (+ 6 mM KCl in medium) to stimulate network activity, we did not detect any significant changes in AcD cell numbers (Fig. 2C, KCl: 16.4% ± 11.0%, cntr: 25.3% ± 22.2%, *P*=0.2309, paired *t*-test, *n*=10).

We used two chemical approaches to reduce network activity by inhibiting NMDA receptors. The application of magnesium (MgSO_4_, +10 mM) blocks NMDA receptors, thus inhibiting synaptic transmission and plasticity ^23^. We found that adding 10 mM MgSO_4_ for seven days did not significantly change the number of AcD neurons, although the values indicate a tendency for a decrease of AcD cell propensity (Fig. 2D, MgSO_4_: 9.9% ± 6.9%, cntr: 22.6% ± 12.6%, *P*=0.0520, paired *t*-test, *n*=10). When presynaptic NMDA receptors were blocked with 1 µM ifenprodil, which dominantly inhibits GluN2B receptors and to a limited degree GluN1/NR2A-containing receptors ^24^, we did not observe any effect on the number of AcD neurons (Fig. 2E, ifenprodil: 13.0% ± 10.6%, cntr: 14.8% ± 24%, *P*=0.8346, paired *t*-test, *n*=6). In summary, bicuculline treatment increased the number of AcD neurons in the treated OTC hemispheres compared to control.

### Neuronal activity modulates AcD development *in vivo*

Our *in vitro* data demonstrated that the axon onset in cortical neurons can be a dynamic feature, transitioning between somatic and dendritic locations. This type of plasticity can be modulated by pharmacologically induced alterations in network activity. However, the question remains whether these morphological remodeling events can occur *in vivo*. There are several critical periods during the development of S1BF during which the network undergoes significant changes in neuronal excitability, synaptogenesis, and balance between excitation and inhibition ^25,26^. We hypothesized that an increase or reduction in sensory input during these critical periods modulates the direction of the preferred AcD transformation in S1BF neurons. Several studies have found that early whisker-specific sensory experiences significantly affect the subcellular anatomy of neurons in the barrel cortex during development ^17,27^. Sensory deprivation through whisker trimming significantly reduces neuronal connections in the deprived cortex ^28^, whereas sensory enrichment increases excitatory activity ^29^. To study the effect of sensory deprivation and activation on axonal onset morphology, we unilaterally trimmed whiskers and compared the deprived hemispheres to the intact/stimulated hemispheres in the same mouse, thereby minimizing inter-mouse variability. We varied the lengths and onsets of the experiments to capture certain critical windows during development (Fig. 3A). Enriched environments (EE) with novel objects introduced daily were incorporated to boost exploration, thus facilitating the imbalance between cut and intact whisker fields.

**Figure 3:**
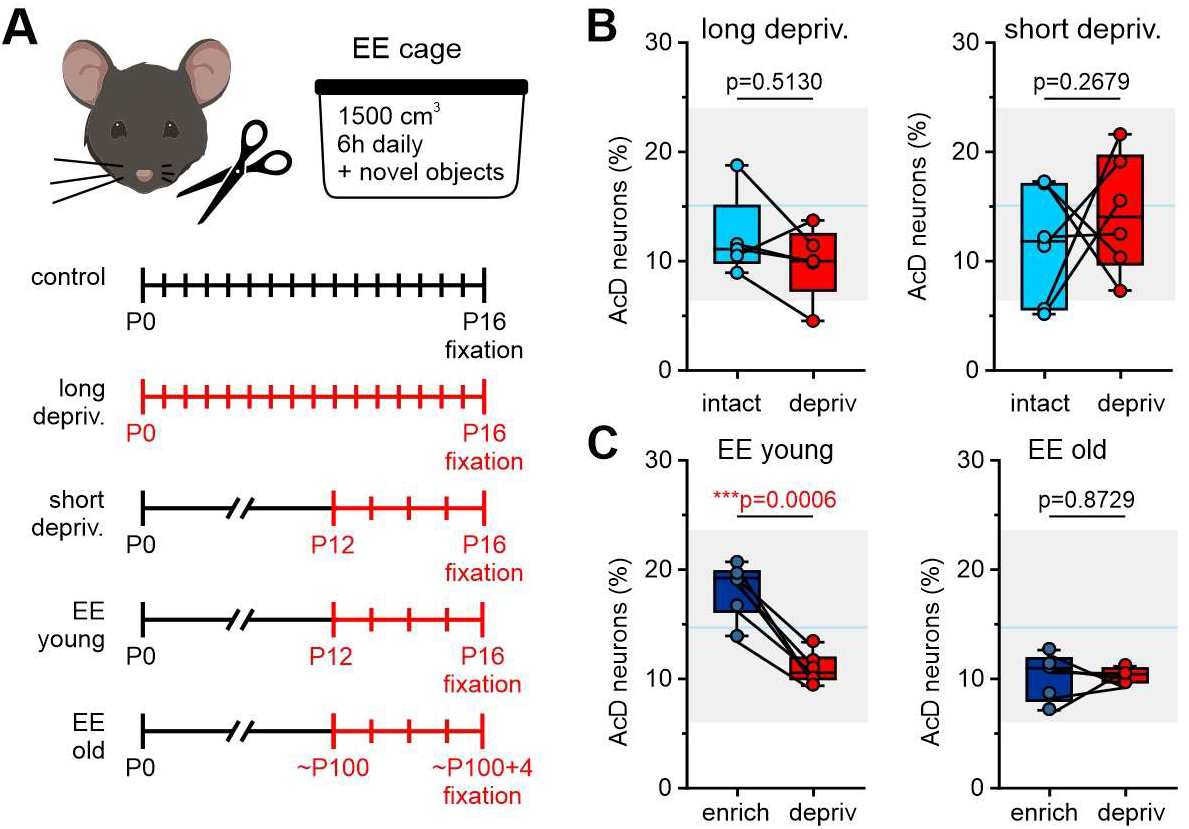
Unilateral whisker deprivation and exploration create asymmetry in AcD cell numbers between both hemispheres. (A) Experimental design for unilateral sensory deprivation. Whiskers were cut daily starting from P0, P12, or < P100 for either 17 or 5 days. During enriched environment experiments (EE), mice were allowed to explore a larger cage containing various beddings, and novel objects with different textures for 6 h daily. (B) The percentage of AcD neurons in the deprived and intact hemispheres remained unaffected during short-term (left, *n*=6) and long-term whisker deprivation (right; *n*=5; paired *t*-test). (C) When whisker deprivation was combined with EE for 5 days in young animals (EE young), we found an asymmetry between AcD cell numbers between the treated and untreated hemispheres (left panel, *P*=0.0006, paired *t*-test, *n*=6). In contrast, the number of AcD neurons was not significantly different when the procedure was performed in older mice (>P100) (left panel, paired *t*-test, *n*=5).

In the long deprivation paradigm, mice were unilaterally trimmed daily from the day of birth (P0) to P16 (Fig. 3A). This approach aimed to examine the enduring effect of sensory deprivation on axonal onset morphology. We hypothesized that an extended period of sensory deprivation might lead to a profound change in the number of AcD neurons. We found that long-term unilateral whisker deprivation had no significant impact on the number of AcD neurons between the deprived and intact S1BF (Fig. 3B; intact: 12.2% ± 3.8%, deprived: 9.9% ± 3.4%, *P*=0.2679, paired *t*-test, *n*=5).

Long-term changes in sensory input lead to rapid network adaptation that could mask the initial plasticity of axon onsets. Thus, we reduced the duration of whisker deprivation to five days. The whisker deprivation window was selected from P12, the start of exploration and active whisking behavior ^25^ to P16 (Fig. 3A). We found that short deprivation caused no change in AcD cell number (Fig. 3B, intact: 11.5% ± 5.3%, deprived:14.4% ± 5.4%, *P*=0.5130, paired *t*-test, *n*=6).

Although the unilateral removal of whiskers is assumed to increase the use of the contralateral/intact side ^30^, the potential for exploration is generally limited in standard housing cages. Thus, we boosted exploration and the use of intact whiskers by exposing the mice to EE for 6 h daily from P12 to P16. At each day of this period, we introduced new objects with novel textures that could be explored by the uncut whiskers. After five days of exploration, we found that cells in the deprived hemisphere contained significantly lower numbers of AcD neurons than those in the enriched hemisphere (Fig. 3C, 10.5% vs. 17.7% of all Thy1-GFP-positive neurons, *P*=0.0006, paired *t*-test, *n*=6). Plasticity in AcD morphology might be favored or restricted to early developmental windows. We thus added an older cohort of mice (>P100) that underwent five days of unilateral whisker trimming, again paired with EE. Overall, the AcD percentage in mature mice was lower than that in juvenile mice. However, there was no discernible difference in the proportion of AcD neurons between the stimulated and deprived hemispheres (Fig. 3C, enriched: 9.7% ± 2.2%, deprived: 9.9% ± 0.7%, *P*=0.8729, paired *t*-test, *n*=5). It is conceivable that the experience of our behavioral paradigm alone has an impact on the number of AcD cells. We thus measured the abundance of AcD cells in mice that were kept in their home cage until P16 without any experimental manipulation. We noted that the variation in the number of AcD neurons between animals decreased when they explored the EE in both deprived and enriched hemispheres as well as in old and young mice (Supplementary Fig.3; *P*=0.0006, Barlett’s test).

In conclusion, a strong imbalance in barrel cortex inputs (unilateral whisker cutting and EE) causes an asymmetric distribution of AcD cells between both hemispheres in juvenile mice.

## Discussion

Neuronal plasticity has been studied extensively for decades and our current understanding centers on the idea that both classical Hebbian as well as homeostatic and other physiological mechanisms are multicomponent events involving significant morphological remodeling of neuronal compartments (spines, dendrites, axons), and (sub)cellular components such as ion channels, receptors or the underlying transcriptome ^31,32^. Morphologically, these types of plasticity include and are not limited to the formation, elimination, and morphological remodeling of synapses, dendritic spines, entire dendrites and axonal compartments such as boutons (reviewed in ^33-36^). A key driving force seems to be neuronal activity, which has been shown to shape network maturation across multiple sensory systems during development, but also later in life (reviewed in ^37,38^).

What could be the benefit of large-scale morphological plasticity as observed in our study? In the hippocampus, AcD neurons offer a unique mode of information processing and network integration. Inputs at AcD branches are more potent, circumvent perisomatic inhibition, and boost network participation during network oscillations such as sharp-wave ripples ^12,13^, an essential component of memory consolidation ^13,39^. Less is known about the functional role of AcD neurons in the somatosensory cortex. Similar to ripples in the hippocampus, this area features ultrafast network oscillations in response to strong, synchronous thalamocortical input, assumed to increase the bandwidth for encoding and transmitting sensory information ^40^. Precise sensory information processing requires fast neuronal response times ^41,42^, and AcD morphology could facilitate the strength and timing of back-propagating action potentials in cortical neurons ^14^. The plasticity of axon onset might therefore optimize the processing of whisker-related inputs and enhance the precision and efficiency of sensory encoding. Neurons activated by particularly salient input patterns may develop AcD morphology to boost detection and optimize response times.

Is large-scale morphological plasticity a mechanism contributing to network homeostasis? We suggest that the modulation of axon onset could provide a mechanism to reconcile Hebbian plasticity with neuronal homeostasis. Strong excitatory input leads to long-term potentiation of respective synapses ^43^ and might thereby trigger a feedback loop that increases firing and network participation of the respective neuron. This can be tolerated to some degree, but mean firing frequencies tend to gravitate towards an optimal mean for individual cells and even more so on the network level. However, if active cells shift their axon onset in response to activity, the efficiency of most dendritic inputs would be reduced while the AcD branch becomes electrotonically privileged ^12^. This would allow the neuron to remain a healthy and meaningful player in the network while sharpening the encoding profile.

Recent studies provide concepts of compartmentalized dendritic plasticity, according to which dendrites receive synaptic inputs from different subcortical and intracortical pathways organized according to their laminar target ^44,45^. In this context, basal dendrites largely receive sensory input, thus playing a distinct role in feedforward information processing ^46^, while the apical and somatic dendritic compartments contribute to sensory encoding, since they receive mostly feedback input ^47-49^. This segregates functional compartments within a single neuron and the authors argue that this architecture provides cortical neurons with a spatial (and temporal?) information-separating mechanism ^44^. Transition of this cortical neuron basal dendrite to or from AcD to nonAcD would then strongly impact neuronal behavior and in combination with other modes of activity-dependent plasticity at the same cell, modulate the computational power of single neurons.

Here, we suggest that the transition in axon onset location as well as its modulation by network activity constitutes an additional mode of morphological plasticity, keeping in mind that we only observed such transitions in real-time in a subset of cells (17 of 98 in total). These transitions are likely governed by different subcellular processes, the precise nature of which remain to be identified. How do differentiated neurons achieve large-scale morphological plasticity? At this point, we can only speculate, but numerous studies have shown that the neuronal cytoskeleton adapts and alter its phenotype to accommodate a single neuron’s needs during events of plasticity (reviewed in ^50^), and we therefore propose that neuronal activity triggers cytoskeletal modulators that mediate a rearrangement from nonAcD to AcD morphology and *vice versa*.

Actin and microtubules are likely involved in large-scale plasticity due to their dynamic nature. While axonal microtubules are usually more stable than those in dendrites ^51-53^, single microtubules can contain both labile and stable domains ^54^, which could be available for large-scale plasticity. Interestingly, “mixed” forms of neurites, which are neither fully dendritic nor axonal in their ultrastructural architecture have been known for decades. The pioneering electron microscopic work on the neuronal cytoskeleton performed by Peters and colleagues described neurons with a shared root, in which the basal dendrite and axon emerge from a common stem. This shared root features characteristics of dendritic fine structure as well as fasciculated microtubules typical for the axon hillock ^2^. In fact, the axon hillock ultrastructure in these neurons as well as in AcD cells exhibits the dense undercoating typical for the AIS, which here begins right after the actual axon branches off this dendritic stem ^2^. How the basal cell pole retains the ability to modulate morphologically will have to addressed in future studies.

In summary, our study unveils a remarkable dynamism of axon onset in layer V cortical neurons in mouse barrel cortex. Our findings provide insights into an underexplored range of morphological parameters potentially available for optimizing and adapting neuronal responses during events of homeostatic plasticity.

### Limitations of the study and outlook

The live observation of axon onset remodeling was performed in OTC, an established *in vitro* culture system with some limitations. While the cytoarchitecture in OTC often remains intact ^55^, the system itself is one of wide-spread axotomy, which in turn can have an effect on (sub)cellular remodeling ^56^. Live-imaging *in vivo* would be the next step to better understand the observed dynamic remodeling mechanisms. Mechanistically, there are a number of questions that should be addressed in future studies, especially in light of recent advances in super resolution microscopy that have shed light on the intricate internal structure of dendrites and axons and can also be applied in live-cell imaging ^50^. How is the cytoskeleton reconfigured at the ultrastructural level when a somatic axon shifts towards a new dendrite? Is the new dendritic stem a thin elongation of the somatic envelope? Or is it feasible to assume that single neurons indeed remodel their entire basal cell pole to shift axon onset, and with it AIS position, when network changes occur? Whatever the mechanism, our data underscore the concept of multicomponent neuronal plasticity, adding to the growing list of morphological parameters that neurons can employ to compensate, adjust and optimize their role in an ever-changing network.

## Acknowledgements

We are indebted to Jan Maximilian Janssen for his introduction to data analysis with the Imaris software, and to Petra Wahle for her continued intellectual support and fruitful discussions of developmental neuroscience topics in general. Our work was funded and supported by the Deutsche Forschungsgemeinschaft, DFG (EN 1240/2-1 to M.E.), the DFG Walter Benjamin Programme (PN 458054460 to C.T.), and intramural funds provided by the Medical Faculty of Johannes Kepler University Linz (to S.M. and M.E.).

## Author contributions

M.E. conceptualized and directed the research. N.L., C.T., and M.E. conceived the experiments. N.L. acquired all data and N.L., S.M., and C.T. analyzed the data. N.L. drafted the original version of the manuscript. N.L., C.T. and M.E. revised and edited the manuscript. All authors read and approved the final version of the manuscript.

## Material and Methods

### Resource availability

Further information and requests for resources and reagents should be directed to and will be fulfilled by the lead contact, Maren Engelhardt.

### Materials availability

There were no new materials generated in the course of this study.

### Data and code availability

All primary data and analysis files will be uploaded to the JKU data management repository and made available upon request. This paper does not report original code. Any additional information required to reanalyze the data reported in this work is available from the lead contact upon request.

### Animals

All animal procedures were carried out in compliance with the guidelines of the Animal Care and Use Committee of the Mannheim Medical Faculty, Heidelberg University, and were approved by the state Baden-Württemberg under the EU guidelines (35-9185/G-119/20). In this study, mixed-gender mice from a transgenic mouse line expressing green fluorescence protein (GFP) under the Thy1 promoter (Tg/Thy1-EGFP) MJrs/J provided by Jackson Laboratory Maine, USA ^57^ were utilized. Mice had access to water and food *ad libitum* and were kept on a regular 12/12 h light/dark cycle at a temperature of 22 ± 2 °C and relative humidity of 45-65%. The offspring were separated from their mothers on postnatal day (P) 28. Experiments were performed between birth and P16, or after P100 on mixed gender background. We did not expect or test for sex related differences.

### Preparation of organotypic slice cultures

Organotypic slice cultures (OTC) were prepared according to previously published methods ^4^. Animals aged P5 were decapitated and their brains placed in ice-cold preparation medium (94,1% Minimum Essential Medium (MEM), 1% GlutaMAX, 2.4% HEPES, 1% D(+)-Glucose (45%), 1% Penicillin/Streptomycin, pH adjusted to 7.4 with 1 M NaOH (0.4% of total volume); all components from Gibco/Thermo Fisher Scientific, Waltham, USA except HEPES from Lonza, Verviers, Belgium and glucose, Pen/Strep from Sigma/Merck, Darmstadt, Germany). Coronal sections were obtained using a vibratome (Microm HM 650V with a cooling chamber, Microm CU65, Thermo Scientific). The cerebellum was removed and the posterior part of the brain was glued onto a vibratome stage. An agarose block (1% dissolved in PBS) was placed on the ventral side of the brain to improve stability during slicing. The slice thickness was set at 275 µm. The hemispheres were separated and transferred onto semipermeable Millicell cell culture inserts (Millipore, 0.4 µm pores, diameter 30 mm, hydrophilic PTFE) and placed in a cooled 6-well plate (Sarstedt AG & Co.KG, Nürnbrecht, Germany). Each well contained 1 ml of preparation medium. After dissecting the brain, Millicell inserts were transferred under sterile conditions into a new sterile 6-well plate, and 37°C warm cell culture growth medium was supplied to each well (42% MEM, 25% Basal Medium Eagle (BME), 25% normal horse serum (NHS), 1% GlutaMAX, 2.5% HEPES, 1.5% D(+)-Glucose (45%), 2% sodium bicarbonate (7.5%), 1% Penicillin/Streptomycin, pH adjusted to 7.3 with 1 M NaOH (0.04% of total volume); all components from Gibco/Thermo Fisher Scientific, Waltham, USA except except HEPES and sodium biocarbonate from Lonza, Verviers, Belgium and glucose, Pen/Strep from Sigma/Merck, Darmstadt, Germany). OTC were cultured in an incubator (Hera Cell 150, Thermo Electron Corporation) at 35°C and 5% CO_2_. Three days after slicing, the growth culture medium was completely exchanged (1 ml) with either control growth culture medium or growth culture medium containing one of the treatments (15 µM bicuculline methiodide, 10 mM MgSO_4_, 6 mM KCl, 1 µM ifenprodile hemitartrate). One hemisphere was treated with one of the above-mentioned reagents, whereas the contralateral hemisphere served as a control (1 µM DMSO). To control for hemispherical differences, treated and untreated sides were alternated. Every second day, half of the medium (500 µl) was replaced.

### Live cell imaging

OTC were generated as described above from the Thy1-GFP mouse line to perform long-term live-cell imaging ^58^. Thy1-GFP positive cells in treated and control OTC were imaged from DIV3 to DIV10 every 24 h. OTC were placed in single sterile tissue culture dishes (Falcon, 35 mm, Easy-Grip Style Cell Culture Dish, Cat. No. 353001, Thermo Fisher Scientific, Waltham, USA). For live-image acquisition, a C2 Nikon confocal microscope (Nikon Instruments, laser line: 488 nm) with a 60x objective (water immersion, numerical aperture of 1.0, CFI Apo 60XW NIR) was used. Images were acquired at 1024 × 1024 pixels in stacks with an acquisition step size of 0.5 µm. For optimal image quality, laser intensity and exposure time were adjusted based on the intensity of the labeled cells. To further enhance the intensity, the pinhole was opened, but never exceeded 2 AU. The imaging process required submerged conditions. We therefore added a column of sterile PBS (37°C warm) onto the slice, which was immediately removed after image acquisition. The OTC was then placed back in the incubator until the next imaging session. Image acquisition lasted for a maximum of 30 minutes per OTC. To avoid contamination, the objective was cleaned with 70% ethanol in water before and after imaging the individual OTC.

### Sensory deprivation/activation in vivo

To manipulate the activity in primary somatosensory, barrel field (S1BF), whiskers were trimmed unilaterally and daily either from P0 to P16, P12 to P16, or for five consecutive days in mice >P100 (see Fig. 3A). Whisker trimming was performed under a binocular microscope by using a curved scissor. Trimming in pups was performed without anesthesia, since young animals showed no signs of distress and could be easily handled. Adult mice (>P100) were briefly anesthetized with isoflurane before trimming to minimize stress during the trimming procedure.

For the sensory activation study, mice with unilaterally trimmed whiskers were placed in an enriched environment (EE) cage for 6 h daily over a period of five consecutive days (see Fig. 3A). The EE cage was significantly larger than the home cage (floor space of home cage 542 cm^3^; cage type 1285L, Tecniplast Germany GmbH), EE cage: 1,500 cm^3^; cage type 1500U, Tecniplast Germany GmbH)), contained different beddings (diverse barks in different sizes, moss, pine cones, different sizes of pebbles, cellulose paper), and novel objects with different textures (wooden bridge, fabric hammock, cardboard tubes, self-rolled bubble wrap tunnel, wooden sticks, and marbles). Novel objects were added daily to EE cages to avoid habituation. The cages were maintained in the dark. One hour prior to exposure to EE, mice were placed in their home cage in a darkened room to adjust to their surroundings.

Unilateral whisker trimming allowed us to compare the putative effects of sensory manipulations within individual mice; thus, the sensory-deprived hemisphere was compared to the hemisphere with intact whisker projection. Up to 5-6 brains were analyzed per group. Per brain and hemisphere, approximately 100 Thy1-GFP neurons in S1BF were analyzed.

### Tissue fixation

Animals P16 and older were put under deep anesthesia with ketamine (120 mg/kg per body weight; Bela-Pharm GmbH & Co., KG, Vechta, Germany) and xylazine (16 mg/kg per body weight; Serumwerk, Bernburg AG, Bernburg, Germany) and transcardially perfused. First, mice were exsanguinated with ice-cold 0.9% NaCl (B. Braun Melsungen AG, Melsungen, Germany) and subsequently fixed with 2 % PFA at room temperature for 5 minutes. Brains were removed from the skull and cryoprotected in 30 % sucrose solution overnight at 4 °C. The following day, the cerebellum was removed and the brain was placed in a Tissue-Tek Cryomold filled with the cryoprotectant Tissue-Tek (both from Sakura Finetek Europe, aan den Rjin, Netherlands). A container filled with isopentane was pre-cooled in liquid nitrogen. Once the isopentane reached the desired low temperature, the cryomold containing the brain was carefully placed in this container and snap-frozen. These samples were stored at -20 °C. Sectioning was performed using a cryostat (Microm, HM 550, Thermo Scientific, Waltham, USA). The tissue block was trimmed to the desired region. Slices of S1BF were dissected coronally from the region of interest at 80 µm thickness and post-fixed in 2 % PFA for 15 min free-floating. The slices were then transferred to 1x PBS and in the dark before staining.

OTC were fixed in 2% PFA for 20 minutes at 4 °C and stored in 1x PBS at 4°C for up to two days until staining was performed.

### Immunofluorescence

Slices were incubated in blocking buffer solution (1% bovine serum albumin, 0.2 % fish skin gelatin, 0.1 % Triton, or 1 % Triton in 1x PBS; BSA from Gibco, Thermo Fisher Scientific, Waltham, USA; FSG from Sigma/Merck, Darmstadt, Germany) for at least 1 h in the dark to block non-specific binding sites and remove background staining. Subsequently, the sections were incubated overnight with the primary antibody solution at 4 °C (blocking buffer solution contained primary antibodies with the following dilution: ch α GFP 1:2000, Abcam; rb α βIV-spectrin 1:1000, self-made ^4^). Subsequently, the slices were washed 3 time with 1x PBS for 10 minutes to remove any remaining unbound primary antibodies. After washing, the slices were incubated in the secondary antibody solution for at least 2 h (all secondary antibodies were diluted 1:1000 in the blocking buffer solution). This was followed by another washing step of 3 × 10 minutes with 1x PBS to remove the redundant antibodies. The slices were then moved onto a cover slide (Epredia, Breda, Netherlands) and dried for approximately 10 minutes. Finally, the slices were immersed in mountain medium (Roti-Mount FluorCare, Carl Roth) with an antifading effect, and covered with a coverslip (Carl Roth GmbH & Co. KG). All slides were dried overnight before confocal microscopy.

In addition, negative controls consisted of the exact procedure, but under omission of the primary antibodies. Secondary antibodies regardless of the species they were directed against did not produce any signals under these conditions.

### Image acquisition

Confocal imaging was performed using a C2 Nikon confocal microscope (Nikon Instruments, laser lines: 488, 543, and 642 nm) with a 60x objective (oil immersion, numerical aperture of 1.4). Images were acquired at 1024 × 1024 pixels in stacks up to 40 µm, with an acquisition step size of 0.5 µm. For optimal acquisition, the exposure intensity was monitored and laser settings were initially set such that the exposure intensity for each laser was normally distributed, and too many oversaturated pixels were avoided.

### Cell classification criteria

Measurements of soma diameter, stem dendrite length, and stem dendrite diameter were performed on confocal imaging stacks in Fiji ^59^, using the segment line tool. Classification was blinded to the treatment groups and underwent a double verification process according to the 4-eyes principle. AcD neurons were defined as cells in which the axon with its AIS branched off from a dendrite. The AcD stem dendrite separating somatic envelope and axonal arbor had to be at least 2.5 µm long and thinner than 20% of the soma diameter. NonAcD cells were defined as (i) having their axon emanating either directly from the soma or (ii) not meeting the criteria for AcD cells. Axon origins at the apical dendrite were classified as AcD, if the distance from the axon to the soma was greater than the diameter of the respective segment. For 3D reconstruction of cell morphology, the surface rendering function of Imaris (v 9.9, Oxford Instruments, Abingdon, UK) was used.

### Statistical Analysis

Statistical tests were performed and graphs were plotted using GraphPad Prism 9 (GraphPad Software, Inc.). AcD cell numbers were captured as ratio between AcD cells and total number of analyzed cells for each hemisphere. Paired or unpaired *t*-test was used for parametric comparisons between the two groups, paired Wilcoxon test for nonparametric comparisons. Boxplots indicate the median, 25% and 75% percentiles (boxes) and 1.5 IQR (Tukey) as well as individual values. *P*-values and number of samples are stated in each figure legend (\**P*<0.05, ** *P*<0.01, *** *P*<0.001).

### Data illustration

Data plots were created in GraphPad Prism and assembled with schematic illustrations in Corel Draw X16 (Corel Corporation, Ottawa, Canada). Fluorescence images were enhanced for contrast using Adobe Photoshop (Adobe Inc., San Jose, CA, USA).

## Acknowledgements

We are indebted to Jan Maximilian Janssen for his introduction to data analysis with the Imaris software.

