## Supplementary Material for "Axon onset remodeling in response to network activity in mouse cortical neurons" (from another version of this preprint: version 1, 2023-08-01)

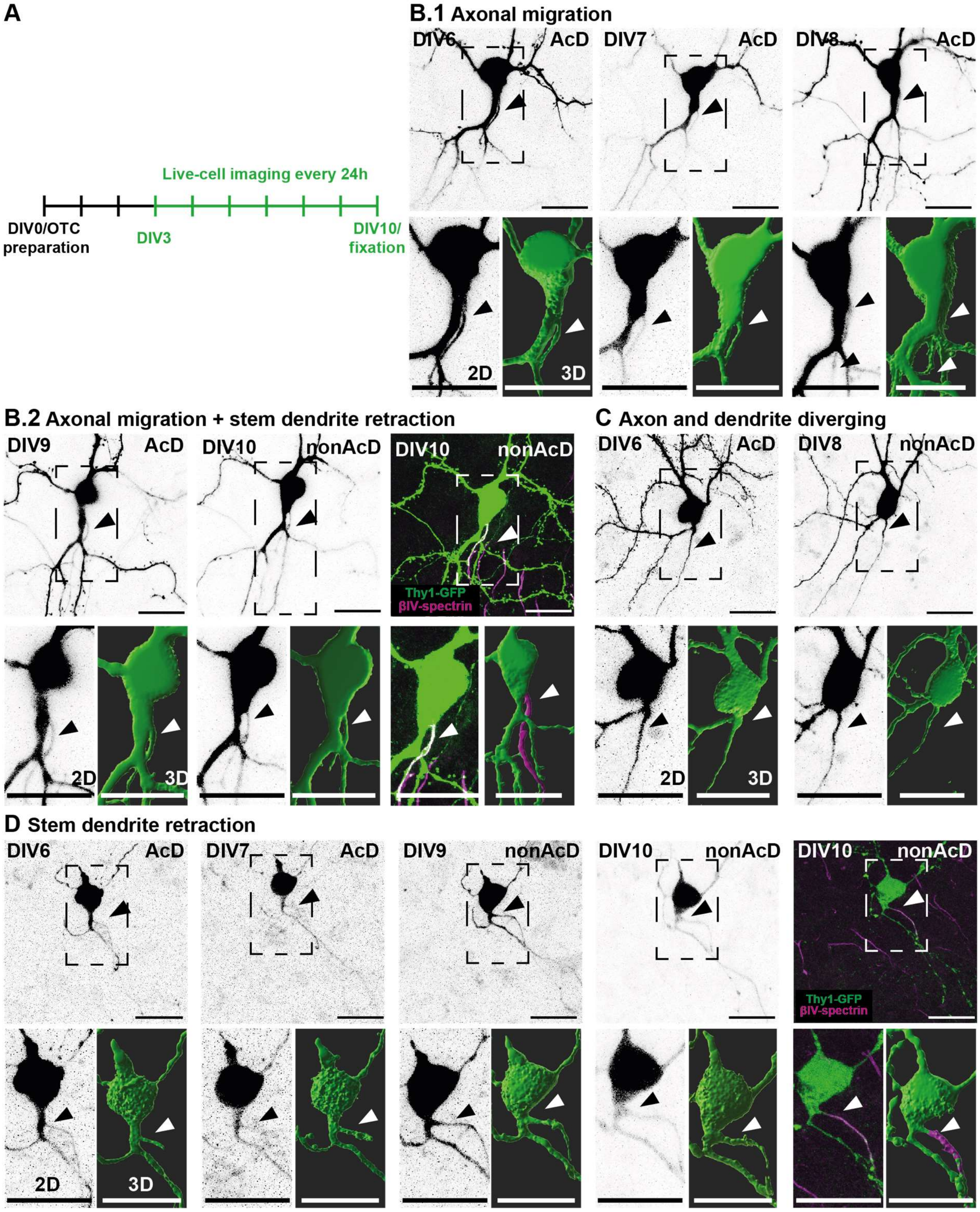


**Supplementary Figure 1: Transition from the AcD to the nonAcD state**. (A) Experimental design. OTCs were prepared from P5 old Thy1-GFP pups. Live-imaging series was conducted from DIV3 to DIV10 every 24h. At DIV10, OTCs were fixed and the AIS was immunolabeled (βIV-spectrin, magenta). (B) The same cell underwent two different pathways for AcD to nonAcD conversion within five consecutive days. B.1: Axon dislocation along the apical dendrite until the axon originated close to the soma. B.2: Axon dislocation accompanied by stem dendrite retraction. (C) Another pathway is axon and dendrite divergence, which was observed within three days. (D) Stem dendrite retraction alone can transform an initial AcD to a nonAcD neuron. Arrowheads indicate the axon origin. Scale bar = 25 μm.

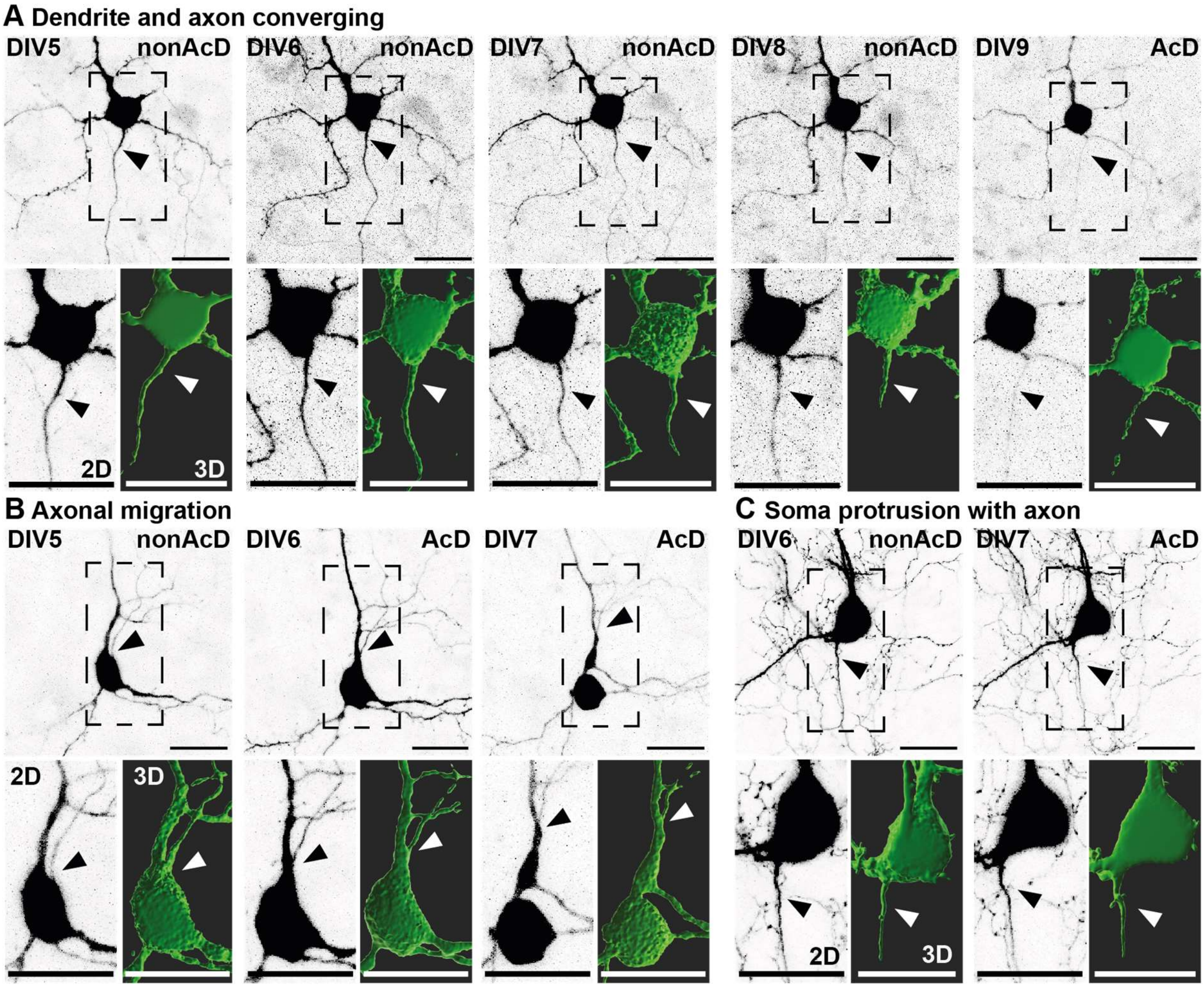


**Supplementary Figure 2: Transition from nonAcD to AcD states**. (A) A basal dendrite converged with an axon, thereby transforming from an initial nonAcD to an AcD neuron within five consecutive days. (B) Axon dislocation along the apical dendrite until the axon originated distal to the soma. (C) Creation of a stem dendrite by a protruding somatic envelope, thus relocating the axon to a more distal position. Arrowheads indicate the axon origin. Scale bar = 25 μm.

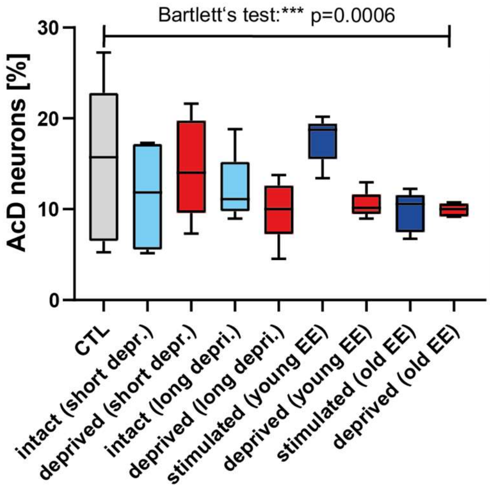


| Treatment | P-value t-test | P-value F-test |
|---|---|---|
| **intact (short depriv.) vs. CTL** | 0.4520 | 0.294 |
| **deprived (short depriv.) vs. CTL** | 0.9197 | 0.3131 |
| **intact (long depriv.) vs. CTL** | 0.5532 | 0.1347 |
| **deprived (long depriv.) vs. CTL** | 0.2766 | 0.0910 |
| **stimulated (EE young) vs. CTL** | 0.4598 | * |
| **deprived (EE young) vs. CTL** | 0.2587 | ** |
| **stimulated (EE old) vs. CTL** | 0.2423 | * |
| **deprived (EE old) vs. CTL** | 0.2476 | *** |

**Supplementary Figure 3: Variance in AcD neurons.** Figure depicts boxplots of the proportion of AcD neurons in the hemispheres of behavioral paradigms applied *in vivo*. P16 control mice were bilaterally unaffected. The bar at the top represents the *P*-value of Bartlett's test in ordinary one-way ANOVA. This result indicated that the variance in the percentage of AcD neurons differed between the behavioral paradigms. Comparing every directly or indirectly affected hemisphere to hemispheres of control mice with regard to AcD percentage showed that the variance in mice that were placed in an EE cage was significantly lower than that in control mice (statistical results depicted in table).

**Supplementary Table 1: Statistical results of the data presented in the study**

| Figure | Group | N | 25% quart. | median | 75% quart. | Mean | SD | SEM | Distr. | p-value | Test |
|---|---|---|---|---|---|---|---|---|---|---|---|
| **Figure 2C** | bicu | 16 | 19.49 | 31.01 | 44.09 | 32.9 | 16.42 | 4.105 | norm | | |
| | CTL | 16 | 2.778 | 17.42 | 20 | 14.58 | 10.17 | 2.517 | norm | 0,0013 | paired *t*-test |
| **Figure 2D** | KCl | 10 | 7.5 | 15.48 | 25 | 16.36 | 11.04 | 3.49 | norm | | |
| | CTL | 10 | 12.64 | 19.44 | 34.37 | 25.34 | 22.2 | 7.02 | norm | 0,2309 | paired *t*-test |
| **Figure 2E** | $MgSO_4$ | 10 | 5 | 10.26 | 14.88 | 9.862 | 6.935 | 2.193 | norm | | |
| | CTL | 10 | 16.67 | 21.54 | 26.44 | 22.62 | 12.6 | 3.986 | norm | 0,052 | paired *t*-test |
| **Figure 2F** | ifenpr | 6 | 0 | 16.52 | 21.25 | 13.01 | 10.64 | 4.342 | norm | | |
| | CTL | 6 | 0 | 0 | 38.89 | 14.81 | 24 | 9.799 | not norm | 0,875 | paired Wilcoxon test |
| **Figure 3B - short depr.** | intact | 6 | 5.538 | 11.84 | 17.18 | 11.49 | 5.297 | 2.162 | norm | | |
| | depr | 6 | 9.588 | 14.03 | 19.77 | 14.41 | 5.409 | 2.208 | norm | 0,513 | paired *t*-test |
| **Figure 3B - long depr.** | intact | 5 | 9.75 | 11.11 | 15.2 | 12.2 | 3.832 | 1.71 | norm | | |
| | depr | 5 | 7.228 | 10 | 12.6 | 9.933 | 3.388 | 1.515 | norm | 0,2679 | paired *t*-test |
| **Figure 3B - young EE** | stim | 6 | 15.52 | 18.73 | 19.44 | 17.74 | 2.494 | 1.018 | norm | | |
| | depr | 6 | 9.45 | 10.14 | 11.64 | 10.5 | 1.434 | 0.5854 | norm | 0,0006 | paired *t*-test |
| **Figure 3B - old EE** | stim | 5 | 7.475 | 10.57 | 11.57 | 9.731 | 2.217 | 0.9913 | norm | | |
| | depr | 5 | 9.185 | 10 | 10.64 | 9.93 | 0.7328 | 0.3277 | norm | 0,8729 | paired *t*-test |

*Bicu = bicuculline; CTL = Control; depr. = deprivation; Distr. = distribution; EE = enriched environment; ifen = ifenprodil; norm = normal distribution; not norm = not normal distribution; quart. = quartile; stim = stimulated*
